# Cardiac myosin binding protein-C palmitoylation is associated with increased myofilament affinity, reduced myofilament Ca^2+^ sensitivity and is increased in ischaemic heart failure

**DOI:** 10.1101/2022.06.21.496992

**Authors:** Alice Main, Gregory N. Milburn, Rajvi M. N. Balesar, Aileen C. Rankin, Godfrey L. Smith, Kenneth S. Campbell, George S. Baillie, Jolanda van der Velden, William Fuller

## Abstract

Cardiac myosin binding protein-C (cMyBP-C) is an essential regulator of cardiac contractility through its interactions with the thick and thin filament. cMyBP-C is heavily influenced by post-translational modifications, including phosphorylation which improves cardiac inotropy and lusitropy, and S-glutathionylation, which impairs phosphorylation and is increased in heart failure. Palmitoylation is an essential cysteine modification that regulates the activity of cardiac ion channels and soluble proteins, however, its relevance to myofilament proteins has not been investigated. In the present study, we purified palmitoylated proteins from ventricular cardiomyocytes and identified that cardiac actin, myosin and cMyBP-C are palmitoylated. The palmitoylated form of cMyBP-C was more resistant to salt extraction from the myofilament lattice than the non-palmitoylated form. Isometric tension measurements suggest c-MyBP-C palmitoylation reduces myofilament Ca^2+^ sensitivity, with no change to maximum force or passive tension. Importantly, cMyBP-C palmitoylation levels are reduced at the site of injury in a rabbit model of heart failure but increased in samples from patients with ischaemic heart failure. Identification of cMyBP-C palmitoylation site revealed S-glutathionylated cysteines C635 and C651 are required for cMyBP-C palmitoylation, suggesting an interplay between the modifications at these sites. We conclude that structural and contractile proteins within the myofilament lattice are palmitoylated, with important functional consequences for cardiac contractile performance.

## Introduction

Cardiac myosin binding protein-C (cMyBP-C) is a sarcomeric accessory protein that has emerged as an essential regulator of cardiac contractility, predominantly through its interactions with the thick and thin filaments (Moss, Fitzsimons and Ralphe, 2015). Three isoforms exist in vertebrate striated tissue, with cMyBP-C exclusively expressed in the cardiac tissue, localised to the C-zone of the sarcomere through interactions of its C-terminal tail domains (C7-C10) with the myosin backbone and titin (Weber *et al.*, 1993; Fougerousse *et al.*, 1998; Flashman, Watkins and Redwood, 2007; Tonino *et al.*, 2019) As such, the N-terminal domains extend from the myosin backbone where they transiently interact with both myosin and the thin filaments (Flashman *et al.*, 2008; Rahmanseresht *et al.*, 2021). Although the exact molecular mechanisms are still unclear, cMyBP-C has been shown to regulate the availability of force generating myosin heads, likely through its interactions with the S1, S2 and regulatory light chain (RLC) components and therefore regulates the rate of force development and force generation of the myofilament (McNamara *et al.*, 2016, 2017, 2019; Sarkar *et al.*, 2020). cMyBP-C also regulates myofilament calcium (Ca^2+^) sensitivity through interactions with the thin filament, including actin, where it binds to reduce rotational flexibility and sliding speed, and tropomyosin, whereby it maintains tropomyosin in its high Ca^2+^ state at low Ca^2+^ concentrations to increase myofilament Ca^2+^ sensitivity and prolong relaxation (Mun *et al.*, 2014; Walcott, Docken and Harris, 2015; Risi *et al.*, 2018; Bunch *et al.*, 2019). Whilst the C0-C2 domains are important in these interactions, the central domains have been less studied, although there is emerging evidence for their role in cMyBP-C stability, structural conformation, protein-protein interactions and post-translational modifications (Gautel *et al.*, 1995; Previs *et al.*, 2016; Doh *et al.*, 2022; Ponnam and Kampourakis, 2022).

Importantly, whilst cMyBP-C can be viewed as a negative regulator of contractility, it is heavily influenced by an increasingly developing profile of post-translational modifications that modulate its function (Main, Fuller and Baillie, 2020). In particular, at least 3 key phosphorylation sites are found in the M-domain (Ser273, Ser282 and Ser302) and cMyBP-C is highly phosphorylated in healthy myocardium (Sadayappan *et al.*, 2005, 2006; Barefield and Sadayappan, 2010; Kumar *et al.*, 2020). Protein kinase A (PKA) mediated phosphorylation reduces the cMyBP-C-myosin interaction, increasing the number of force-generating heads and improving cardiac inotropy. Additionally, phosphorylated cMyBP-C shows reduced binding to actin, improving cardiac lusitropy (Kensler, Shaffer and Harris, 2011; Bunch *et al.*, 2019; McNamara *et al.*, 2019). Importantly, cMyBP-C phosphorylation is frequently attenuated in ischaemic heart failure and age-related cardiac dysfunction, and its preservation in animal models is cardioprotective (Sadayappan *et al.*, 2006; El-Armouche *et al.*, 2007; Jacques *et al.*, 2008; Anand *et al.*, 2018; Rosas *et al.*, 2019). As such, there has been a concerted effort to characterise the post-translational modification profile of cMyBP-C and any links to cardiac disease. This has identified several novel modifications, including cysteine modifications S-nitrosylation and S-glutathionylation, the latter of which is associated with impaired phosphorylation and is increased in human heart failure (Patel, Wilder and John Solaro, 2013; Stathopoulou *et al.*, 2016; Budde *et al.*, 2021).

Palmitoylation is a regulatory modification that has been demonstrated in every class of protein, in every tissue investigated to date, including the myocardium (Essandoh *et al.*, 2020). The modification involves the enzymatic addition of a fatty acid (most commonly palmitate (16C) derived from palmitoyl-CoA) to the thiol of a cysteine residue, catalysed by a group of 23 transmembrane zDHHC-palmitoyl acyltransferases (zDHHC-PATs) and reversed by soluble acylthioesterases (Main and Fuller, 2021). The addition of the hydrophobic fatty acid often leads to targeting or attachment of the substrate to intracellular membranes, although it has also been demonstrated to regulate protein stability, protein-protein interactions and other post-translational modifications (Linder and Deschenes, 2007; Salaun, Greaves and Chamberlain, 2010; Jia *et al.*, 2014). In cardiac tissue, palmitoylation regulates membrane transporters and is required for NCX1 inactivation, phospholemman mediated inhibition of the Na^+^/K^+^ ATPase, conductance of potassium channels and mitochondrial transport (Tian *et al.*, 2010; Tulloch *et al.*, 2011; Reilly *et al.*, 2015; Main, Robertson-Gray and Fuller, 2020; Plain *et al.*, 2020; Amanakis *et al.*, 2021). Importantly, palmitoylation has been implicated in the pathophysiology of myocardial infarction, where increased activity of zDHHC5 contributes to massive endocytosis and cardiac damage following reperfusion (Hilgmann *et al.*, 2013; M. J. Lin *et al.*, 2013). Although palmitoylation plays an important regulatory role for several cardiac substrates, its relevance for myofilament proteins has not been investigated. Recent mass spectrometry evidence suggests myofilament proteins undergo palmitoylation (Miles *et al.*, 2021) and in the present study, we identified that myosin, actin and cMyBP-C are palmitoylated in ventricular cardiomyocytes. Upon further investigation, we found that increased cMyBP-C palmitoylation was associated with increased resistance to salt extraction from the myofilament lattice and reduced myofilament Ca^2+^ sensitivity. Importantly, cMyBP-C palmitoylation levels were dysregulated in both rabbit and human heart failure samples. Additionally, we identified two cysteines in the central domains that undergo S-glutathionylation which are required for cMyBP-C palmitoylation, implicating an important interplay between the two modifications.

## Results

### Cardiac actin, myosin and cardiac myosin binding protein-C are palmitoylated in rabbit, adult/neonatal rat and mouse ventricular tissue

We used acyl-resin assisted capture (Acyl-RAC) to purify palmitoylated proteins from ventricular cardiomyocytes isolated from adult rabbit, rat and mouse, as well as homogenised ventricular tissue from neonatal rat hearts. Acyl-RAC captures proteins via cysteines revealed by hydroxylamine cleavage of the thioester bond that links fatty acids to these cysteines after alkylation of ‘unoccupied’ cysteines under denaturing conditions. Acyl-RAC revealed that ~10-15% of myosin, actin and cMyBP-C are all palmitoylated in these tissues, whilst no measurable palmitoylation was detected for tropomyosin, troponin-T (TnT) or troponin-I (TnI). The specificity of acyl-RAC for fatty acylated proteins lies in the selectivity of neutral hydroxylamine for the thioester bond. However, since myofilament proteins may be subject to multiple reversible cysteine post-translational modifications we ruled out non-specific capture in this assay by conducting acyl-RAC assays from cardiac lysate pre-treated with DTT (to reduce cysteine disulfides and nitrosothiols) prior to cysteine alkylation (Supplementary Figure 1). As it is constitutively palmitoylated with a high stoichiometry, Caveolin-3 was included as an acyl-RAC assay control (Figure 1; Howie et al., 2014)

**Figure 1.**
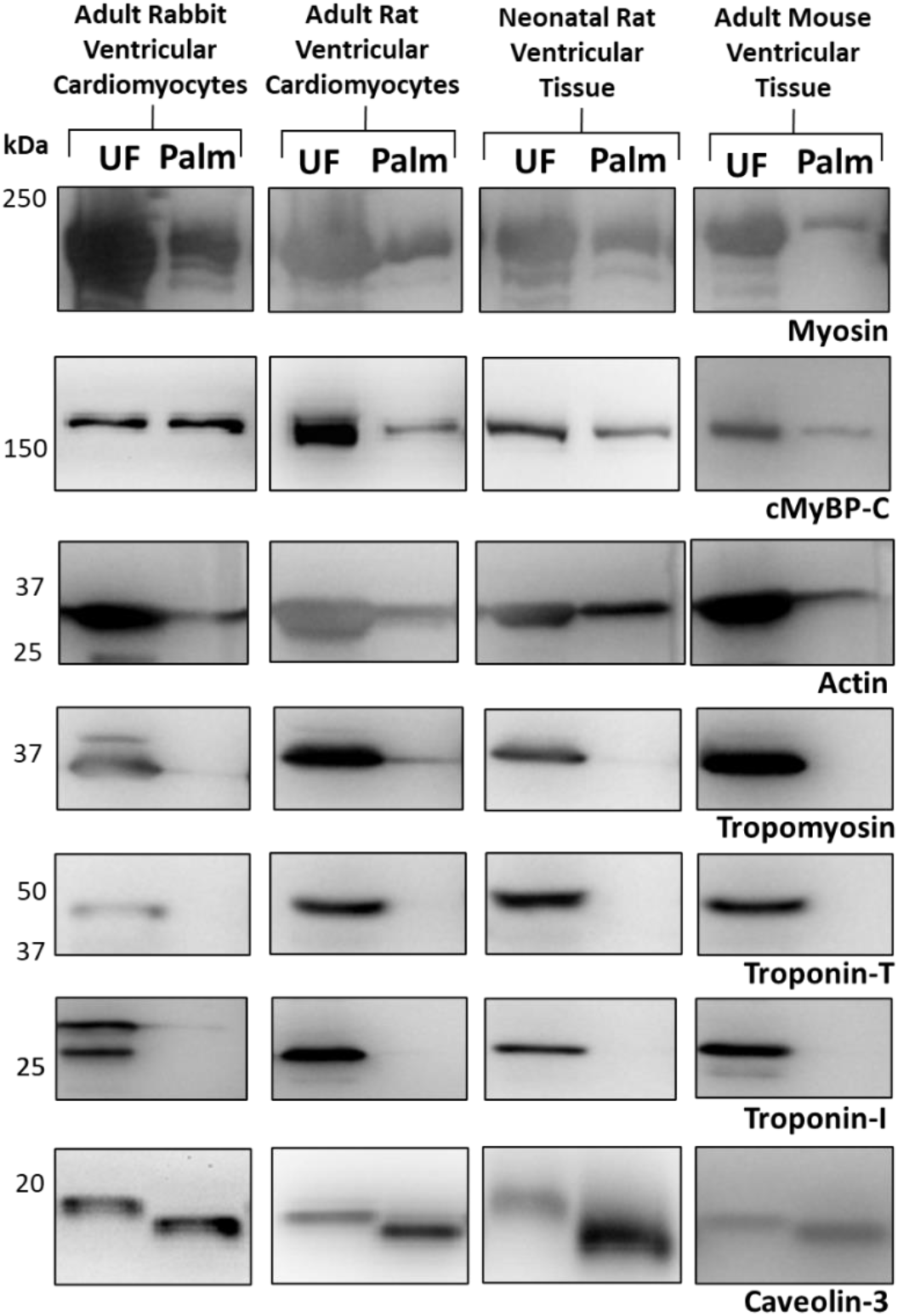
Palmitoylation of thick and thin filament proteins in cardiac tissue. Acyl-resin assisted capture (Acyl-RAC) was used to purify palmitoylated proteins from rabbit ventricular cardiomyocytes (male, 12-week-old), rat ventricular cardiomyocytes (male, 12-week-old), neonatal rat ventricular tissue (male and female, 1-4 days old) and adult mouse ventricular tissue (male, 20 weeks old). Palmitoylation of myosin, cMyBP-C, actin, tropomyosin, troponin-T troponin-I and assay control Caveolin-3 was determined and palmitoylated fraction (Palm) normalised to total protein (UF, unfractionated). Acyl-RAC revealed myosin, actin and cMyBP-C are palmitoylated in all animal model tissues. Representative image of n=3-8.

### Cardiac MyBP-C is palmitoylated in the ventricular myofilament and is resistant to salt extraction

The cardiac isoform of MyBP-C is expressed in cardiac tissue exclusively, but is located in both ventricular, septal and atrial tissue (Sadayappan and de Tombe, 2012; B. Lin *et al.*, 2013). Region-specific variations in cMyBP-C PTMs have never been reported. We purified palmitoylated proteins from the left and right ventricles of the rabbit heart and revealed cMyBP-C may be more palmitoylated in the left ventricle compared to the right (Figure 2A). In ventricular cardiomyocytes, cMyBP-C is anchored to the myofilament predominantly by electrostatic interactions of its C-terminal domains with the LMM portion of the myosin backbone and the giant sarcomeric accessory protein titin (Lee *et al.*, 2015). Titin, along with other cytoskeletal components are insoluble in concentrated salt solutions, whilst ~50% of myofilament proteins are solubilised in 500mM sodium chloride solution (López-Bote, 2017; Xiuping Li *et al.*, 2022). To investigate the subcellular localisation of palmitoylated cMyBP-C and its resistance to salt extraction, myofilaments were isolated from rabbit left ventricular cardiomyocytes and the myofilament pellet further fractionated using 500mM sodium chloride (Figure 2B). The majority of myofilament-localised cMyBP-C was found to be soluble in sodium chloride, with a smaller fraction being insoluble, however when we investigated cMyBP-C palmitoylation in these fractions, this insoluble fraction had a higher level of palmitoylation compared to the soluble fraction (Figure 2C).

**Figure 2.**
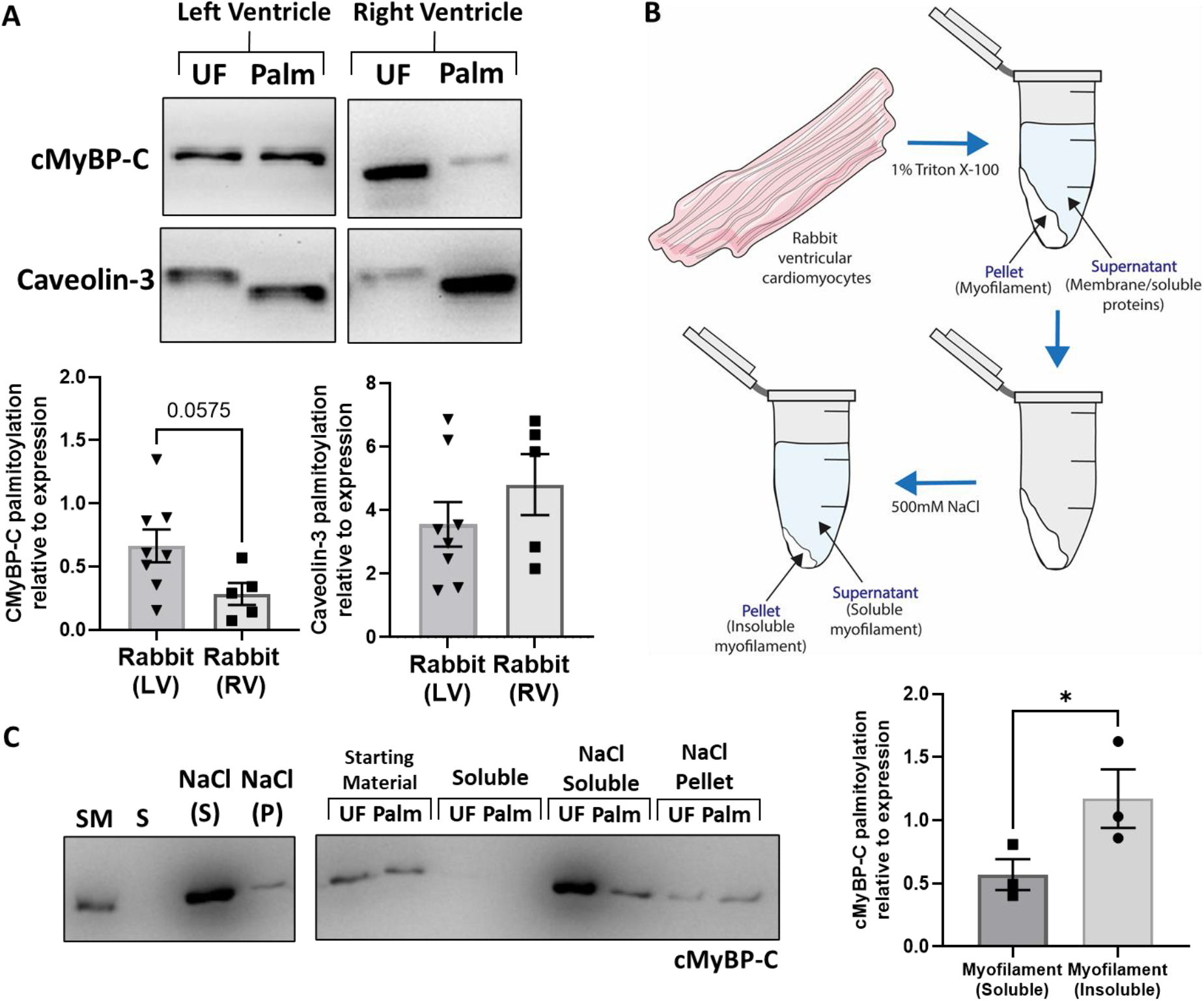
Localisation of cMyBP-C palmitoylation in ventricular cardiomyocytes. **A) CMyBP-C may be more palmitoylated in the left ventricle compared to the right.** Cardiomyocytes were isolated from the left ventricule (LV) and right ventricule (RV) of the adult rabbit heart. Palmitoylation of cMyBP-C and assay control Caveolin-3 was determined by Acyl-Resin Assisted Capture (AcylRAC) and the palmitoylated fraction (palm) normalised to total protein (UF, unfractionated). Statistical comparison between left and right ventricle made via an unpaired student’s t-test. Data is mean ±S.E.M. **B) Graphical schematic of myofilament fractionation.** Rabbit ventricular cardiomyocytes were homogenised in F60 buffer with 1% Triton X-100 and centrifugation used to separate soluble (cytosolic and membrane fraction) and insoluble (myofilament) fractions. The myofilament fraction was further fractionated into myofilament proteins soluble in 500mM NaCl (soluble myofilament) and those insoluble in 500mM NaCl (insoluble myofilament). **C) The palmitoylated form of cMyBP-C is resistant to salt extraction.** During the myofilanent fractionation, a sample was taken at each stage and demonstrated that cMyBP-C as being entirely localised in the myofilament. The majority of cMyBP-C was found to be soluble in NaCl, with a smaller fraction insoluble, however this insoluble fraction of cMyBP-C had a greater proportion of palmitoylated MyBP-C than the soluble fraction (*p<0.05). Palmitoylation of cMyBP-C and assay control Caveolin-3 was then determined by Acyl-Resin Assisted Capture (AcylRAC) and palmitoylated fraction (palm) normalised to total protein (UF, unfractionated). Statistical comparisons made via an unpaired student’s t-test. Data is mean ±S.E.M.

### Palmitoyl CoA treatment increases cMyBP-C palmitoylation and decreases myofilament calcium sensitivity

Interest in palmitoylation as a dynamic modification has been hampered by the assumption that it was a non-enzymatic modification that relied on local concentration of acyl CoA. This is because substrates, including the DHHC-PATs themselves, are auto-palmitoylated on cysteines in the presence of palmitoyl-CoA (Rana, Lee and Banerjee, 2018). Whilst enzymatic regulation of palmitoylation has now been established, tools to induce auto-palmitoylation remain a powerful means to study the effect of increasing substrate palmitoylation. To investigate more directly cMyBP-C palmitoylation, isolated myofilaments were treated with palmitoyl-CoA which spontaneously palmitoylates cysteines and this action led to a significant increase in overall cMyBP-C palmitoylation. To determine the functional consequence for the myofilament of MyBP-C palmitoylation, measurements of passive force (F_passive_) and active force were made at varied calcium concentrations using a muscle mechanics set up. Whilst there was no significant effect on the force generated in maximally activating calcium (pCa4.5, F_max_), there was a significant reduction in the force generated by palmitoyl CoA treated myofilaments in response to submaximal calcium (pCa6.0, F_act_) suggesting a reduction in myofilament calcium sensitivity (FigureB). There was no significant effect of palmitoyl CoA treatment on phosphorylation of S282 (Figure 3A) or on myofilament passive force (Supplementary Figure 2).

**Figure 3.**
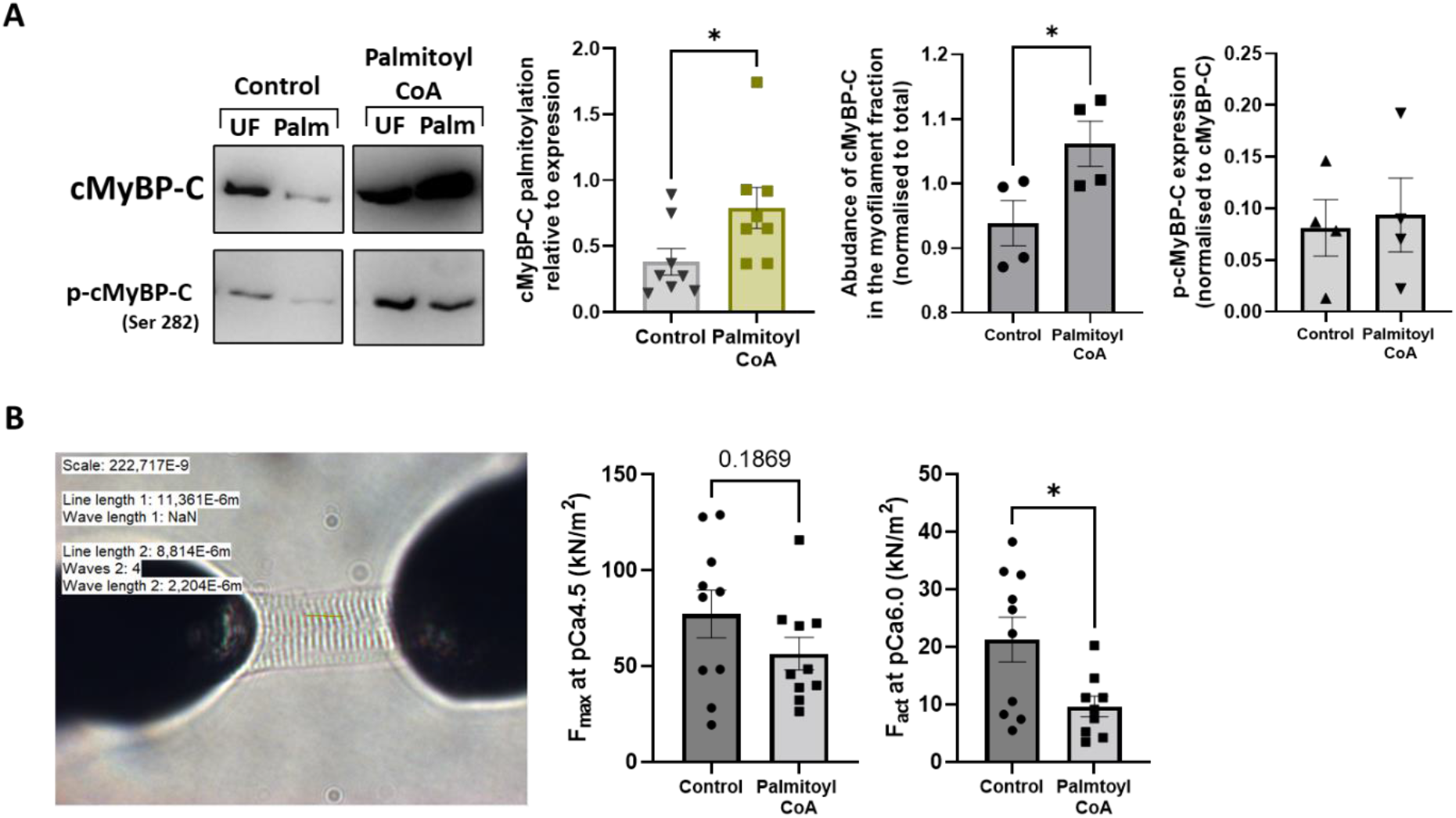
Palmitoyl CoA treatment of isolated myofilaments. The effect of palmitoyl CoA (20μM), which spontaneously palmitoylates solvent exposed cysteines on myofilaments isolated from rabbit ventricular cardiomyocytes. Myofilaments were treated for 30 minutes before centrifugation and collection of the myofilament pellet and supernatant. **A) Palmitoyl CoA treatment of isolated myofilaments increases cMyBP-C palmitoylation and abundance in the myofilament**. Palmitoyl CoA significantly increased cMyBP-C palmitoylation in the myofilament (*p<0.05). Analysis of cMyBP-C abundance revealed that palmitoyl CoA treatment led to a higher abundance (*p<0.05) of cMyBP-C in the myofilament. Palmitoyl CoA treatment had no effect on level of p-cMyBP-C (S282) in the myofilament. Palmitoylation in each fraction was determined by Acyl-Resin Assisted Capture (Acyl-RAC) and total protein (UF, unfractionated) compared to palmitoylated fraction (palm). Data is mean ±S.E.M analysed via an unpaired student’s t-test. **B) Palmitoyl CoA treatment of isolated myofilaments reduces myofilament calcium sensitivity.** Myofilaments were stretched to 2.2μm using a muscle mechanics set up and active force was measured in maximally activating Ca2+ (F_max_ pCa4.5) and submaximal Ca2+ (F_act_ at pCa6.0). Treatment with palmitoyl CoA had no effect on maximal active force in maximally activating Ca2+ but significantly reduced the force generated in response to submaximal Ca2+ (p<0.05). N=9-10 cells from 4 animals. Data is mean ±S.E.M analysed via an unpaired student’s t-test.

### cMyBP-C is palmitoylated at previously identified S-glutathionylation sites C623 and C651

Solvent exposed cysteines that are palmitoylated are often targeted by other cysteine modifications such as glutathionylation, which has been reported to have antagonising effects on protein palmitoylation (Burgoyne et al., 2012; Howie et al., 2013). Glutathionylation of cMyBP-C has been previously reported at C623 and C651 (human sequence) and increased S-glutathionylation is associated with an increased myofilament calcium sensitivity (Patel, Wilder and John Solaro, 2013). As we observed lower force at submaximal calcium with increased cMyBP-C palmitoylation, we investigated whether these sites were also palmitoylated. FLAG-cMyBP-C was exogenously expressed in HEK293 cells where it was palmitoylated and localised to the cytoplasm (FigureA). Mutation of C623 and C651 in full length FLAG-cMyBP-C led to a significant loss in palmitoylation indicating that these cysteines are palmitoylated, albeit not exclusively (FigureB).

### cMyBP-C palmitoylation is reduced in a rabbit model of heart failure and increased in ischaemic human heart failure

Changes in cMyBP-C post-translational modifications including phosphorylation, acetylation and glutathionylation are associated with ischaemic heart failure in animal models and patients, with loss of phosphorylation being most associated with myofilament dysfunction (Sadayappan *et al.*, 2006; Copeland *et al.*, 2010; Govindan *et al.*, 2012; Stathopoulou *et al.*, 2016). To determine the relevance of cMyBP-C palmitoylation in the context of cardiac disease, we investigated a rabbit model of heart failure. Cardiomyocytes were isolated from the left ventricular and septal region of failing and control rabbit hearts, and palmitoylation of cMyBP-C was determined by Acyl-RAC. In the left ventricle, cMyBP-C palmitoylation was significantly reduced in the MI model compared to control, whilst no significant difference was observed in the septal region or in assay control Caveolin-3 (Figure).

To determine whether cMyBP-C palmitoylation is altered in ischaemic heart failure, Acyl-RAC was used to purify palmitoylated proteins from ventricular endocardium samples from ischaemic human heart failure patients and organ donor controls (details in Supplementary Table 1). Analysis of cMyBP-C expression revealed no significant differences, whilst phosphorylation at S282 was reduced although not significantly (p=0.0537). There was a significant increase in cMyBP-C palmitoylation in heart failure samples compared to organ donors (Figure). In contrast, in samples from hypertrophic cardiomyopathy patients there was no significant change in cMyBP-C palmitoylation, regardless of sarcomeric or non-sarcomeric mutation, compared to organ donor controls (Supplementary Figure 1).

## Discussion

In this study, we report for the first time that myofilament proteins actin, myosin and cMyBP-C are post-translationally palmitoylated. With a focus on cMyBP-C palmitoylation, we find that this may be region-specific within the heart and that whilst palmitoylated cMyBP-C is exclusively localised in the myofilament, it is more resistant to salt extraction than non-palmitoylated cMyBP-C. Treatment of myofilaments with palmitoyl CoA enhances cMyBP-C palmitoylation and these myofilaments show reduced myofilament Ca^2+^ sensitivity but no change in maximal force or passive tension. Both cysteines 623 and 651, previously reported to be sites of cMyBP-C glutathionylation, are required for cMyBP-C palmitoylation. In a rabbit model of heart failure, cMyBP-C palmitoylation is significantly reduced in the remodelled myocardium 8 weeks post-MI. In contrast, cMyBP-C palmitoylation is significantly increased in samples from ischaemic heart failure patients compared to organ donors.

### Palmitoylation is a novel post-translational modification of myofilament proteins

Palmitoylation is an essential regulatory modification in the cardiac tissue, but the focus has primarily been on its role in membrane transport and membrane-associated signalling (Jacqueline Howie *et al.*, 2013; Reilly *et al.*, 2015; Gök and Fuller, 2020; Gök *et al.*, 2020; Main and Fuller, 2021). In the present study, Acyl-RAC was used to identify novel palmitoylation of sarcomeric proteins myosin, actin and cMyBP-C in ventricular tissue from rabbit, rat and mouse. Additionally, thin filament-based proteins TnI, TnT and tropomyosin were not found to be palmitoylated in these samples (Figure 1). In line with the data presented, the Tseng group have recently published a study characterising the cardiac palmitoylome using mass spectrometry, identifying more than 400 proteins with cMyBP-C, myosin and actin among them (Miles *et al.*, 2021). Interestingly, the study also identified TnT, TnI and tropomyosin, despite the evidence presented here that they are not palmitoylated in cardiac tissue. These proteins were also identified in our own mass spectrometry analysis of the rat heart, altogether emphasising the importance of further validation of palmitoylation proteins identified in mass spectrometry studies (*Unpublished*, Fuller et al.). The subcellular localisation of DHHCs is based on exogenous HEK293 studies and an understanding the distribution of the enzymes in cardiac tissue specifically would be a valuable tool (Ohno *et al.*, 2006). The palmitoylating enzymes are integral membrane proteins, so proximity to the sarcomere to palmitoylate sarcomeric proteins is difficult to envisage. We therefore don’t rule out non-enzymatic palmitoylation of cMyBP-C occurring through spontaneous reaction with cytosolic acyl-CoA, although the vast majority of palmitoylation is thought to be enzymatic. Future investigations will need to address the subcellular location of these enzymes in the myocardium.

### Palmitoylation of cMyBP-C increases its resistance to salt extraction, reduces calcium sensitivity and occurs on functionally important cysteines

The fraction of cMyBP-C that is palmitoylated is more resistant to salt extraction from the myofilament lattice, a fractionation technique usually employed to break the electrostatic interactions within the myofilament (Figure 2C). This could indicate that the palmitoylated form of cMyBP-C is more tightly associated with the myofilament lattice – possibly as a result of additional hydrophobic interactions mediated by the fatty acid. Alternatively, it is conceivable that palmitoylated cMyBP-C represents a population attached to closely localised membrane structures, potentially aided by the hydrophobic environment created by adding the palmitoyl group(s) (Xiaoling Li *et al.*, 2022).

Given the importance of cMyBP-C PTMs in regulating cardiac contractility, and the difference in force generation and electrophysiological properties between the left and right ventricles, it is surprising there is a lack of study of the extent of PTMs between the anatomical regions (Molina, Heijman and Dobrev, 2016; Pham *et al.*, 2019). We identified modestly higher cMyBP-C palmitoylation in the left ventricle compared to the right, although whether this correlates to the function of the two ventricles remains to be determined (Figure 2A). Notably, our functional analysis indicates enhanced cMyBP-C palmitoylation reduces myofilament Ca^2+^ sensitivity. cMyBP-C palmitoylation had no significant effect on passive force (Supplementary Figure 2) or on the ability of the myofilaments to generate maximal force in saturating Ca^2+^ (pCa4.5), however, there was a significant decreased in the maximal force generated in submaximal Ca^2+^ (pCa6.0) suggesting a loss of Ca2+ sensitivity (Figure 2B). Increased cardiomyocyte fatty acid availability is associated with high fat diets, diabetes and insulin sensitivity (Zhao *et al.*, 2018; Glatz, Luiken and Nabben, 2020) and at a cardiomyocyte level, treatment with high concentrations of palmitate or a mixture of fatty acids was associated with a reduction in peak sarcomere shortening and reduced myofilament calcium sensitivity which we suggest may be mediated in part by increased cMyBP-C palmitoylation (Angin *et al.*, 2012; Zhao *et al.*, 2016).

We identified that both C623 and C651 are required for some, but not all, of cMyBP-C palmitoylation (Figure 4B). The functional importance of these sites in cMyBP-C is already established. Increased S-glutathionylation at C623 and C651 is correlated with increased myofilament Ca^2+^ sensitivity and slower cross-bridge kinetics (Patel, Wilder and John Solaro, 2013). Not only may palmitoylation and glutathionylation compete to modify these cysteines (as has been observed in other substrates (Burgoyne *et al.*, 2012; Jacquieline Howie *et al.*, 2013)), they also induce functionally opposing effects. It is interesting that both these modifications occur in the lesser studied central domains of cMyBP-C, as opposed to the myosin/titin binding C-terminal domains or thick and thin filament regulating N-terminal domains. There is a growing appreciation of the importance of this region of the protein, including potentially in forming bent or hinge conformations of the protein, as an enzyme docking site, in binding of the S1 domain of myosin and as a region of extensive PTM modification (Gautel *et al.*, 1995; Previs *et al.*, 2016; Doh *et al.*, 2022; Ponnam and Kampourakis, 2022). It will be important to determine whether the increased hydrophobicity and myofilament retention as a result of cMyBP-C palmitoylation contributes to or antagonises any of these reported mechanisms.

**Figure 4.**
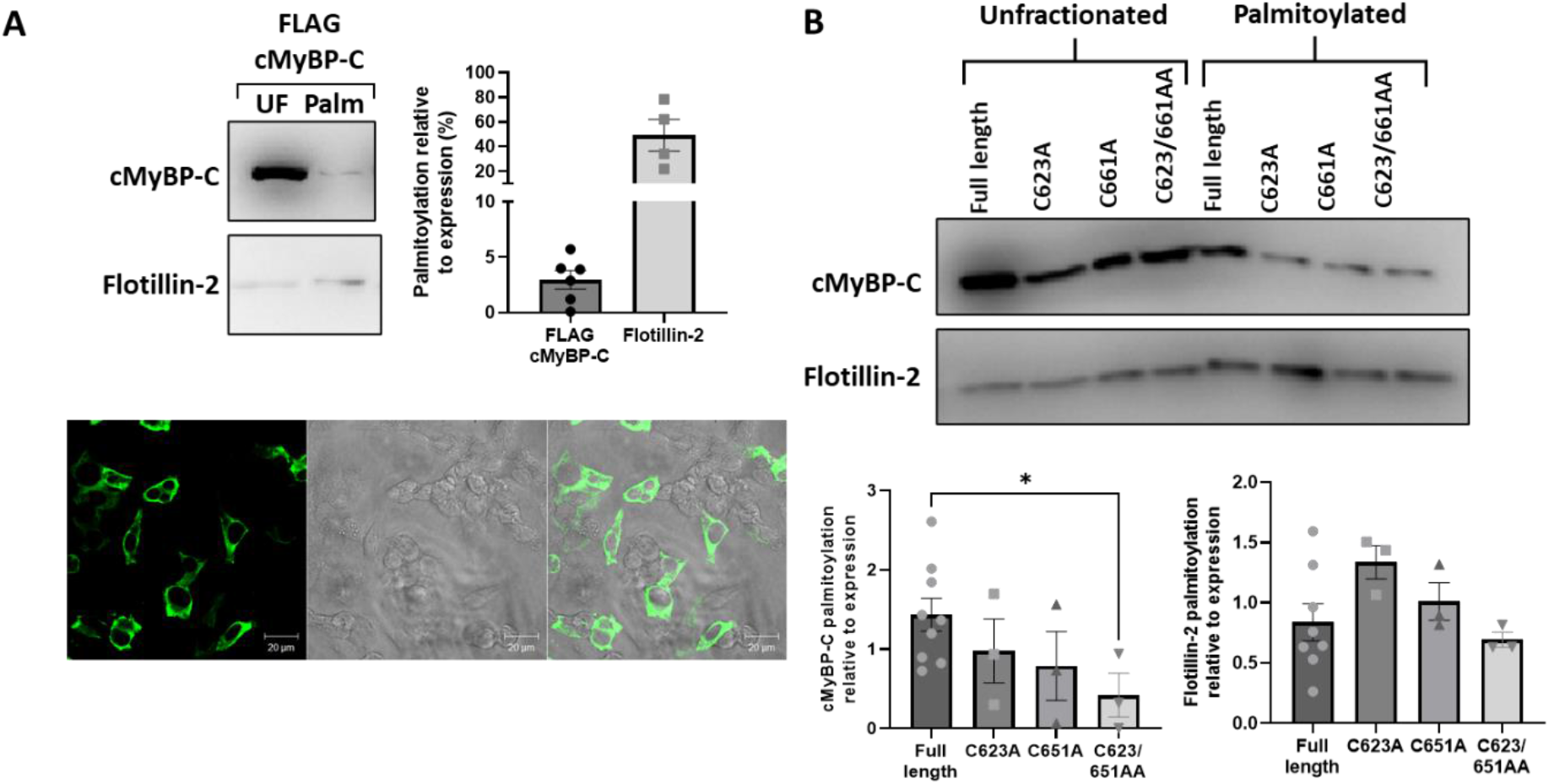
Exogenously expressed FLAG-cMyBP-C palmitoylation. **A) FLAG-cMyBP-C is palmitoylated in HEK293 cells.** Acyl-resin assisted capture (Acyl-RAC) of transiently transfected HEK293 cells with 1μg of FLAG-cMyBP-C revealed cMyBP-C is palmitoylated in these cells. Confocal microscopy of transfected cells revealed that FLAG-cMyBP-C expression is largely cytoplasmic and not clearly localised to any membrane compartment. Scale bar is 50μm. **B) FLAG-cMyBP-C requires C623/651 to be palmitoylated.** Mutation of both C623 and C651 to alanine in FLAG-cMyBP-C results in a loss of palmitoylation (*p<0.05). Palmitoylation of cMyBP-C and assay control Flotillin-2 were determined by Acyl-Resin Assisted Capture (AcylRAC) and palmitoylated fraction (Palm) normalised to total protein (UF, unfractionated). Statistical comparisons made via a One-way ANOVA with a Dunnet’s post-hoc test. Data is mean ±S.E.M.

### Palmitoylation is a therapeutically relevant post-translational modification of cMyBP-C

The disease relevance cMyBP-C PTM regulation, particularly concerning phosphorylation, acetylation and S-glutathionylation, has been demonstrated using animal models of cardiac injury and in samples from patients of HCM and heart failure (Copeland et al., 2010; Govindan, Sarkey, et al., 2012; Sadayappan et al., 2006; Stathopoulou et al., 2016). In this study, cMyBP-C palmitoylation is reduced in the remodelled myocardium 8-weeks post-MI, but only in tissue isolated from the left ventricular region (infarct, peri-infarct and remote), but not the remote septal region (Figure 5). In the LV, the peri-infarct region undergoes the most substantial remodelling and haemodynamic stress, whilst the septal region of the rabbit heart experiences secondary remodelling (Burton *et al.*, 2000; Konstam *et al.*, 2011). This could indicate that cMyBP-C palmitoylation changes are occurring in the area that is most damaged or undergoing the most significant remodelling, including that of T-tubules (Setterberg et al., 2021). However, in contrast, samples obtained from organ donors and ischaemic human HF patients showed reduced cMyBP-C phosphorylation, although not significantly, and an increase in cMyBP-C palmitoylation (Figure 6). Interestingly, HCM caused by *MYBPC3* mutations, other sarcomeric mutations or non-sarcomeric mutations did not result in a significant change in cMyBP-C palmitoylation, suggesting that the changes in palmitoylation are specific to the development of ischaemic heart failure (Supplementary Figure 3). Relatively little is known about the remodelling of palmitoylating enzymes in heart failure of HCM, and it will be important to determine whether this contributes to the contrasting changes observed in animal models and humans. Additionally, there are many confounding factors in human samples that may influence cMyBP-C phosphorylation and palmitoylation including sex, history of cardiac injury and presence of hyperlipidaemia (Supplementary Figure 4, 5 and 6) although the relatively small sample size make statistically relevant comparisons challenging overall.

**Figure 5.**
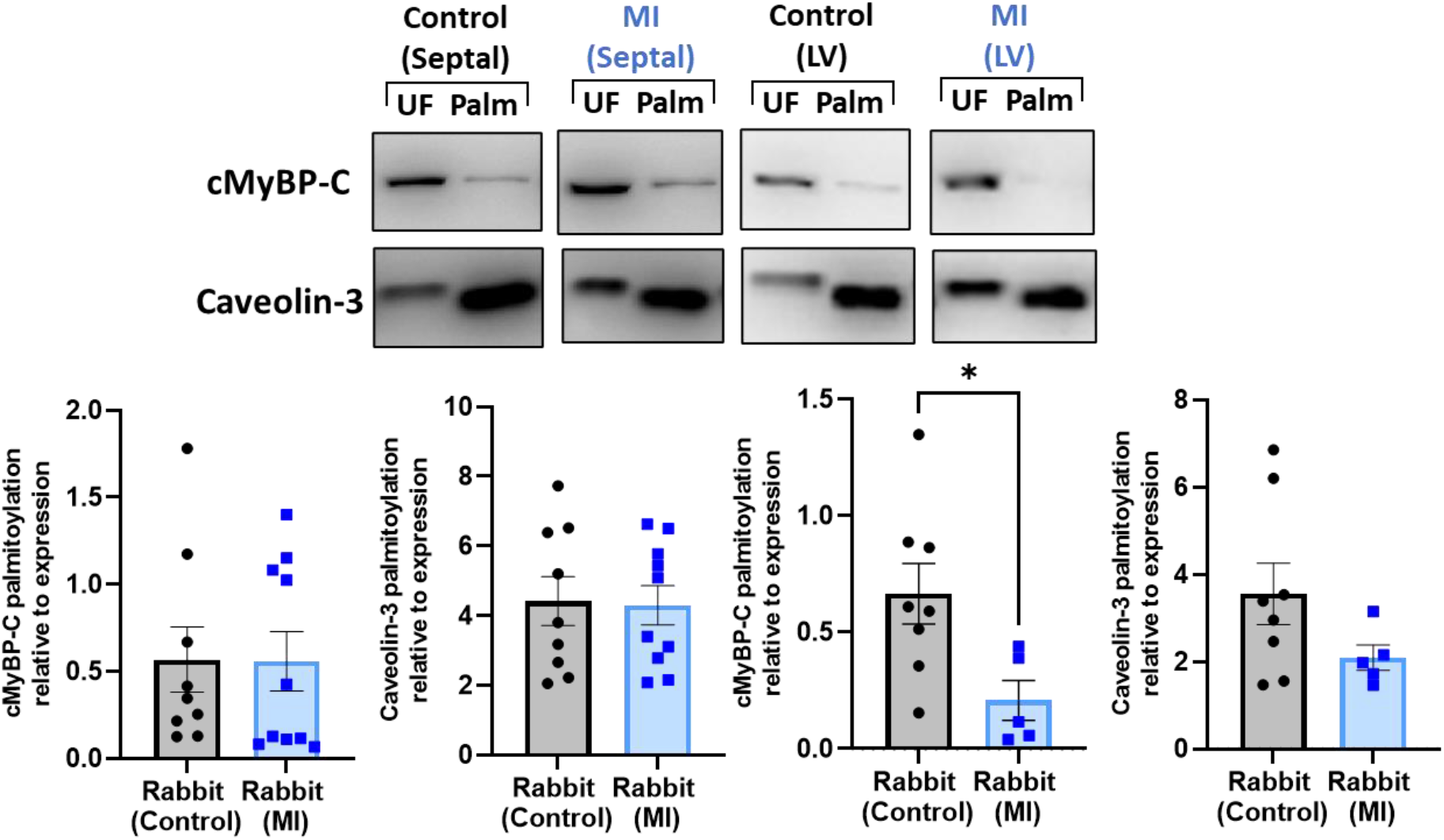
cMyBP-C palmitoylation is reduced in a rabbit model of heart failure in the left ventricular but not septal region. Ventricular cardiomyocytes were isolated from septal and left ventricular (LV) regions of the rabbit heart. Rabbits were either stock/sham at 12/20 weeks of age, or a model of myocardial infarction (MI) whereby the rabbits underwent left anterior descending coronary artery ligation at 12 weeks of age before being maintained for a further 8 weeks. Palmitoylation of cMyBP-C and assay control Caveolin-3 was determined by Acyl-Resin Assisted Capture (Acyl-RAC) and palmitoylated fraction (palm) normalised to total protein (UF, unfractionated). Palmitoylation of cMyBP-C was reduced in cardiomyocytes from the left ventricle, but not septal, region of the rabbit heart following MI (*p<0.05). Data is mean ±S.E.M analysed by a student’s unpaired t-test.

**Figure 6.**
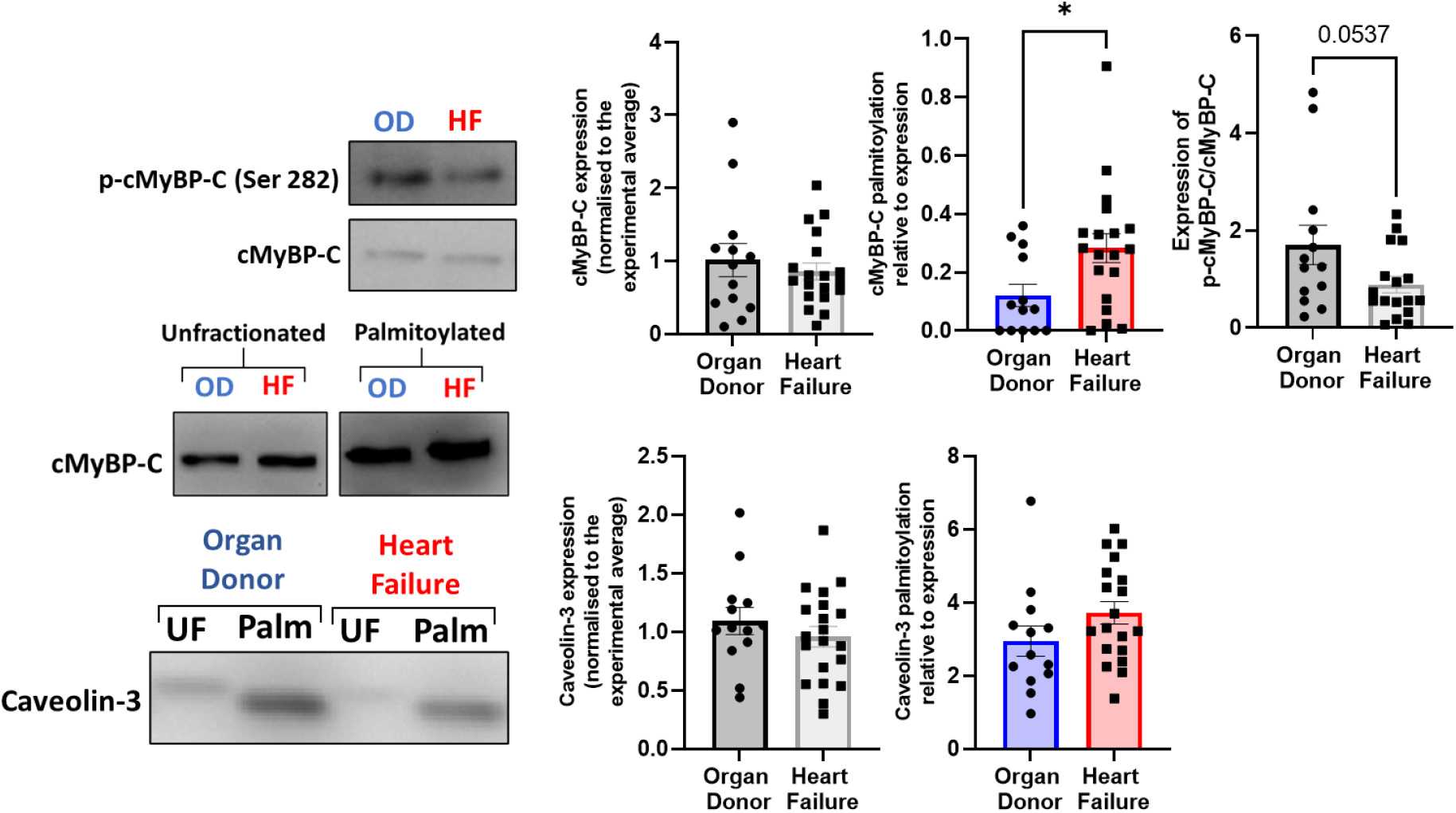
cMyBP-C palmitoylation is increased in samples from ischaemic heart failure patients. Expression and palmitoylation of cMyBP-C and assay control Caveolin-3 was determined by Acyl-Resin Assisted Capture (Acyl-RAC) in ventricular endocardium from organ donor and ischaemic heart failure samples, represented as palmitoylated protein (palm) normalised to total protein (UF, unfractionated). Palmitoylation of cMyBP-C is significantly increased in heart failure samples compared to organ donor (*p<0.05), with no significant change in overall expression and a trending decrease in phosphorylation (p=0.0537). Data is mean ±S.E.M analysed via an unpaired student’s t-test.

### Methodological limitations and future directions

We acknowledge that the functional evidence presented in this investigation does not allow an understanding of the exact role of cMyBP-C palmitoylation, as other sarcomeric proteins may be palmitoylated following incubation with palmitoyl-CoA. Utilising myofilaments from WT and cMyBP-C KO mice would allow the contribution of cMyBP-C palmitoylation to be determined, as has achieved in the study of cMyBP-C S-glutathionylation (Stathopoulou *et al.*, 2016; Cuello *et al.*, 2018). Recently, a novel method has been developed by the Harris lab involving a transgenic mouse in which the C0-C7 domains of cMyBP-C can be “cut” away from isolated myofilaments and replaced rapidly by “pasting” a recombinant version in its place (Napierski *et al.*, 2020). Using this method, a recombinant form of C0-C7 with increased palmitoylation could be introduce and the effect on myofilament parameters determined, with the opportunity for in-vivo study.

Interestingly, whilst palmitoylation was also identified in both skeletal isoforms of MyBP-C to a similar extent of the cardiac isoform (Supplementary Figure 7), C623 is a site unique to cMyBP-C, whilst C651 is only shared with the fast skeletal isoform. This is not uncommon for cMyBP-C modifications, as of the three M-domain phosphorylation sites, only one site exists in the slow skeletal isoform and none in the fast skeletal isoform (McNamara and Sadayappan, 2018). Whilst C623 and C651 may be the predominant sites of cMyBP-C palmitoylation, several of the cysteines within MyBP-C are poorly conserved between isoforms, indicating that the skeletal isoforms may have their own sites that could contribute to the overall similar level of palmitoylation. In that regard, a major a limitation of detecting palmitoylation sites in HEK293 cells is that whilst they can provide a useful system to detect the presence or absence of palmitoylation, they show low levels of palmitoylated cMyBP-C and lack of proper cellular location. In cardiomyocytes, cysteine availability may be dependent on contractile state, and therefore palmitoylated cysteines detected in HEK293 may not be as important for endogenous MyBP-C in cardiomyocytes or skeletal muscle cells (Gross and Lehman, 2013). This has previously been observed for phosphorylation of cMyBP-C, where higher levels were observed in-vivo, including the identification of 9 new sites, compared to in-vitro and ex-vivo studies (Kooij *et al.*, 2013).

## Methods

### Human organ donor and ischaemic heart failure samples

Samples from organ donors (non-failing) and human heart failure patients were obtained from the Gill Cardiovascular Biorepository at the University of Kentucky, of which patients and families of organ donors provided written consent. All procedures were approved by the local IRB and details of the collection procedures have been published previously (Blair *et al.*, 2016). Samples in this study were taken from the ventricular endocardium of each heart with details in Supplementary Table 1.

### Human organ donor and hypertrophic cardiomyopathy samples

Samples from organ donors (non-failing) and hypertrophy cardiomyopathy patients were obtained from the University of Amsterdam Medical Centre, of which patients and families of organ donors provided written consent with collection procedures have been published previously (Schuldt et al., 2021). Samples in this study were taken from the ventricular endocardium of each heart with details in Supplementary Table 2.

### Ethical statement

Animals were handled in accordance with the UK Animals (Scientific Procedures) Act of 1986. All procedures were approved by the UK Home Office (PP7088535) and Glasgow University Ethics Review Committee. The animal research reported here adheres to the ARRIVE and Guide for the Care and Use of Laboratory Animals guidelines.

### Rabbit cardiomyocyte isolation and model of heart failure

Adult rabbit ventricular myocytes (ARVM) were isolated from New Zealand white rabbits weighing between 3-4kg, as previously described. Isolation of adult rabbit ventricular cardiomyocytes (ARVM) was completed as described previously (Kettlewell *et al.*, 2013). Briefly, New Zealand White male rabbits (12 weeks old, ~3-4kg) were euthanised with a terminal dose of sodium pentobarbital (100mg/kg) with heparin (500IU), following which the heart was removed and retrogradely perfused on a Langendorff system. Enzymatic digestion of the tissue using a protease and collagenase solution occurred for ~15 minutes before heart was removed from the system and cut into sections (left atria, right atria, right ventricle, left ventricle and septal regions) and each was finely dissected in Krafte-Brühe solution. The mixture was then triturated and agitated for 10 minutes before filtering and the cell suspension was centrifuged manually for a minute before the pellet was re-suspended in fresh KB. For experiments, cells were stepped up to physiological calcium in modified Krebs-Henseleit solution, initially containing 100μM of CaCl2 and left to settle for before the process was repeated using 200μM, 500μM, 1mM and 1.8mM concentrations of CaCl2. Cells were then snap frozen and kept at −80°C or used for functional experiments.

### Subcellular fractionation of rabbit cardiomyocytes

Isolated cardiomyocytes were harvested in F60 buffer (60mM KCl, 30mM Imidazole, 2mM MgCl_2_•6H_2_O) with 1% Triton X-100 before an analytical sample was taken (unfractionated) and the remaining lysate centrifuged at 14,000G for 5 minutes at 4°C (Kuster *et al.*, 2015). This allows separation of the soluble components of the cell (supernatant) and the myofilaments (pellet). An additional step was taken to resolubilise the myofilament pellet with 500mM sodium chloride followed by centrifugation at 14,000G for 5 minutes at 4°C and separation of the soluble and insoluble fractions.

### Myofilament isolation and pharmacological treatment

To determine the effect of pharmacologically palmitoylating myofilament proteins, myofilaments from adult rabbit cardiomyocytes were isolated and treated before analysis via Acyl-RAC or isometric force experiments. Whilst frozen cells could be used for biochemical measurements, for functional experiments fresh cells were used. In both cases cells were incubated with relaxing buffer (Na2ATP (5.9mM), MgCl2 (6.04mM), EGTA (2mM), KCl (139.6mM), Imidazole (10mM) in dH20, pH 7.4, pCa9.0) containing 0.5% triton and rotated at 4°C for 30 minutes to permeabilise the membrane and reveal the myofilaments. Myofilaments were then centrifuged at 4°C at 1200RPM for 5 minutes before the supernatant was removed and discarded. The resulting myofilament pellet was then washed in relaxing buffer three times with centrifugation to remove any residual triton. For treatment, myofilaments were resuspended in relaxing buffer containing palmitoyl alkyne-coenzyme A (palmitoyl-CoA; 20μM; Cambridge Biosciences, CAY15968) and rotated at 4°C for a further 30 minutes before centrifugation. For isometric force experiments, myofilament pellets were resuspended in a 50% glycerol solution in relaxing buffer and stored at −20°C until use.

### Myofilament passive and active force measurements

Mechanical myofilament measurements were performed using a muscle mechanics workstation as previously described (van der Velden *et al.*, 1999). The system was maintained at 15°C using a water bath and myofilaments viewed using an inverted microscope and a 5X (0.12NA) or a 32X (0.4NA) objective. Skinned myofilaments (preparation described in **Error! Reference source not found.**) were placed in relaxing buffer on a glass coverslip and individual cells were selected based on size and uniformity of striations. Individual myofilaments were then attached to thin, stainless-steel needles with one attached to a piezoelectric motor (Physike Instrumente, Waldbrunn, German) and the other attached to a force traducer (SensoNor, Horten, Norway). Overall cell length, width and sarcomere length was determined by using a CCD video camera and a spatial Fourier transform algorithm. Passive tension of the myofilament (F_passive_) was determined by rapid shortening to 30% of the length of the cell at three different sarcomere lengths ranging from 2μM to 2.2μM (±0.005μM) which has been reported to encompass the working range of the heart (**Error! Reference source not found.**). The depth of the cell was estimated to be 80% of cell width. Active and passive (F_passive_) force of development of the myofilaments at a sarcomere length of 2.2μM was determined in maximally activating solution (pCa4.5, F_max_), followed by submaximal Ca^2+^ (pCa6.0) and then maximal activating solution again. The run down between the first and second activating solutions could then be determined and those with large rundowns (>25%) indicating potential cell detachment/loss of sarcomere length were excluded from analysis.

### Acyl-resin assisted capture

Acyl-resin assisted capture was used to purify palmitoylated proteins in a sample and adapted from a method published previously (Forrester *et al.*, 2011). Cultured and pelleted cells were lysed in the SDS buffer (2.5% SDS, 1mM EDTA, 100mM HEPES pH7.4) containing 1% methyl methanethiosulfonate (MMTS) to methylate free cysteines. Proteins were then precipitated using acetone at with the resulting pellet subsequently washed using 70% acetone to remove excess MMTS. The pellets were then re-suspended in 1% SDS, 1mM EDTA, 100mM HEPES pH 7.5 before a portion of the sample was taken (unfractionated sample, total protein). To the remaining solution, 250mM NH2OH (hydroxylamine (HA), pH 7.5) was added to hydrolyse thioester bonds. Proteins with free cysteines were the purified using thiopropyl sepharose beads (GE Healthcare). Palmitoylation of substrates was determined by quantifying the amount capture relative to unfractionated lysates.

### Immunoblotting

Standard western blotting was carried out using 6-20% gradient gels. The primary antibodies used were as follows: cMyBP-C (1:2000, Santa Cruz Biotechnology, sc-137237), p-cMyBP-C (1:1000, pSer^282^, Enzo Life Sciences, ALX-215-057-R050), myosin heavy chain (1:200, Developmental Studies Hybridoma Bank, MF20), cardiac actin (1:200, Developmental Studies Hybridoma Bank, JLA20), troponin-T (1:200, Developmental Studies Hybridoma Bank, CT3), troponin-I (1:200, Developmental Studies Hybridoma Bank, TI-1), tropomyosin (1:200, Developmental Studies Hybridoma Bank, CH1), slow skeletal-MyBP-C (1:1000, MYBPC1, Caltag Medsystem (ProSci), PSI-6679), fast skeletal-MyBP-C (1:1000, MYBPC2, Caltag Medsystem (ProSci), PSI-5651), FLAG (1:1000, Sigma-Aldrich, G3165), Flotillin-2 (1:1000, BD Biosciences, 610383), Caveolin-3 (1:4000, BD Biosciences, 610420),. The secondary antibodies used were as follows: Rabbit anti-mouse HRP (1:2000, Jackson ImmunoResearch 111-035-144), goat anti-rabbit HRP (1:2000, Jackson ImmunoResearch 315-035-003), goat anti-mouse IgG H&L AlexaFluor® 488 secondary antibody (1:500, Thermofisher A-21085).

### Statistics

Statistical analysis was completed using GraphPad Prism (Version 7; California, USA) and was completed on groups with 3 biological replicates or more. For comparisons in data sets with more than two groups, a one-way analysis of variance (ANOVA) with a Dunnett’s or a Tukey post hoc test was used, with comparisons detailed in the figure legend. For comparisons of two groups an unpaired student’s t-test was used. All samples were analysed for the presence of significant outliers (Grubb’s test). A probability of p<0.05 was considered to be statistically significant.

## Acknowledgements

We acknowledge the patients and families of organ donors who donated cardiac samples for this research.

## Availability of data and material

Further information and requests for resources and reagents should be directed to and will be fulfilled by the corresponding author, William Fuller

## Authors’ contributions

AM: Conceptualization, Investigation, Formal Analysis, Writing – original draft

WF: Conceptualization, Project Administration, Funding Acquisition, Supervision, Writing – review & editing

GB: Funding Acquisition, Supervision and Methodology

JVDV: Resources and methodology

GM, AR, GS, KC: Resources

