## Supplementary Figures for "Cardiac myosin binding protein-C palmitoylation is associated with increased myofilament affinity, reduced myofilament Ca^2+^ sensitivity and is increased in ischaemic heart failure"

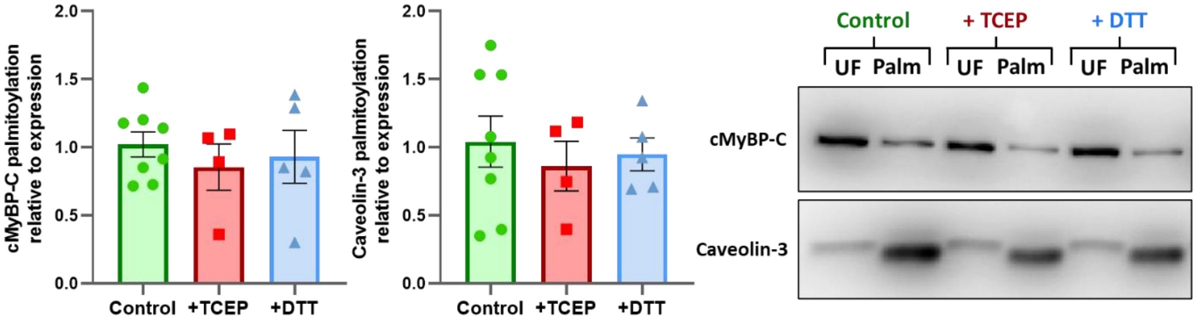


**Supplementary Figure 1. Palmitoylation of cMyBP-C in the presence of Tris(2-carboxyethyl)phosphine (TCEP) and Dithiothreitol (DTT).** Prior to detection of palmitoylated cMyBP-C, rabbit cardiomyocyte lysates were treated for 10 minutes with 10mM DTT or TCEP to reduce disulphides. Palmitoylation of cMyBP-C and assay control Caveolin-3 was then determined by Acyl-Resin Assisted Capture (Acyl-RAC) and palmitoylated fraction (palm) normalised to total protein (UF, unfractionated). There was no significant difference in palmitoylation level with TCEP or DTT addition compared with no pre-treatment (control). Data is mean ±S.E.M analysed via a one-way ANOVA with a Dunnett's post-hoc test.

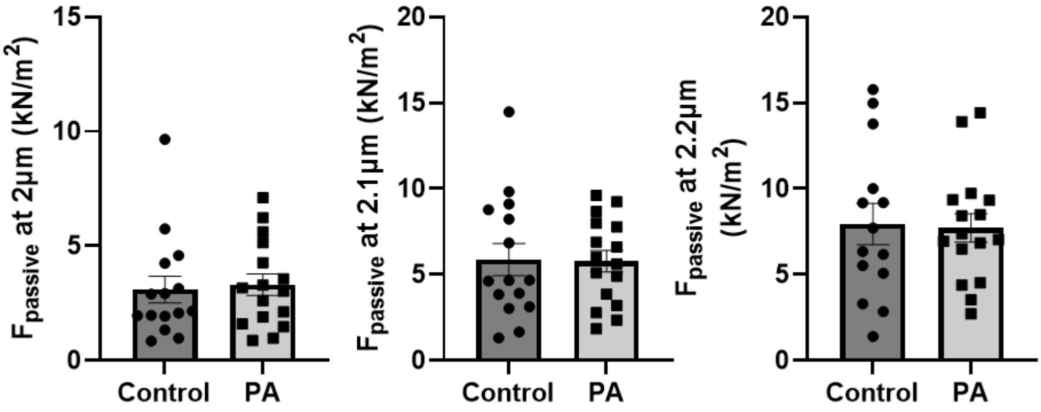


**Supplementary Figure 2. Passive force in palmitoyl CoA treated myofilaments.** Myofilaments were isolated from rabbit ventricular cardiomyocytes and treated with palmitoyl CoA (20µM) before mounting to a muscle mechanics set up. Myofilaments were stretched to 3 different sarcomere lengths (2µm, 2.1µm and 2.2µm) and passive tension was determined by slackening cells by 30% of their length and measuring the difference in force pre and post slacking. Treatment with palmitoyl CoA had no significant effect on passive tension at any sarcomere length. N=15-16 cells from 4 animals. Data is mean ±S.E.M analysed via an unpaired student's t-test.

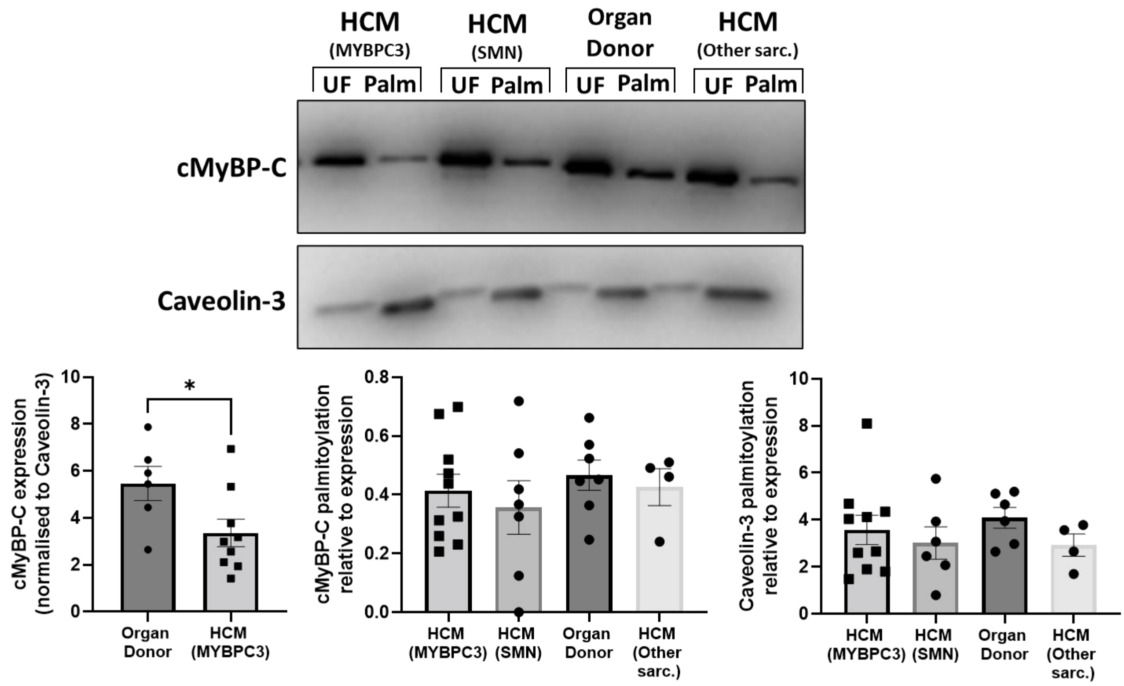


**Supplementary Figure 3. cMyBP-C palmitoylation is unchanged in hypertrophic cardiomyopathy samples.** Palmitoylation of cMyBP-C and assay control Caveolin-3 was determined by Acyl-Resin Assisted Capture (Acyl-RAC) in ventricular tissue from organ donor and hypertrophic cardiomyopathy (HCM) samples with either *MYBPC3* mutations, non-sarcomeric mutations (SMN) or other sarcomeric mutations (*MYH7, MYL2* and *TNNT2*). *MYBPC3* mutations are associated with a loss of cMyBP-C expression resulting from haploinsufficiency (*p<0.05, normalised to Caveolin-3 expression). Despite a change in cMyBP-C expression, there was no significant difference in cMyBP-C palmitoylation suggesting both palmitoylated and unpalmitoylated cMyBP-C are lost at the same rate in HCM. Results are represented as palmitoylated protein (palm) normalised to total protein (UF, unfractionated). Data is mean ±S.E.M analysed via a One-way ANOVA with a Tukey post-hoc test.

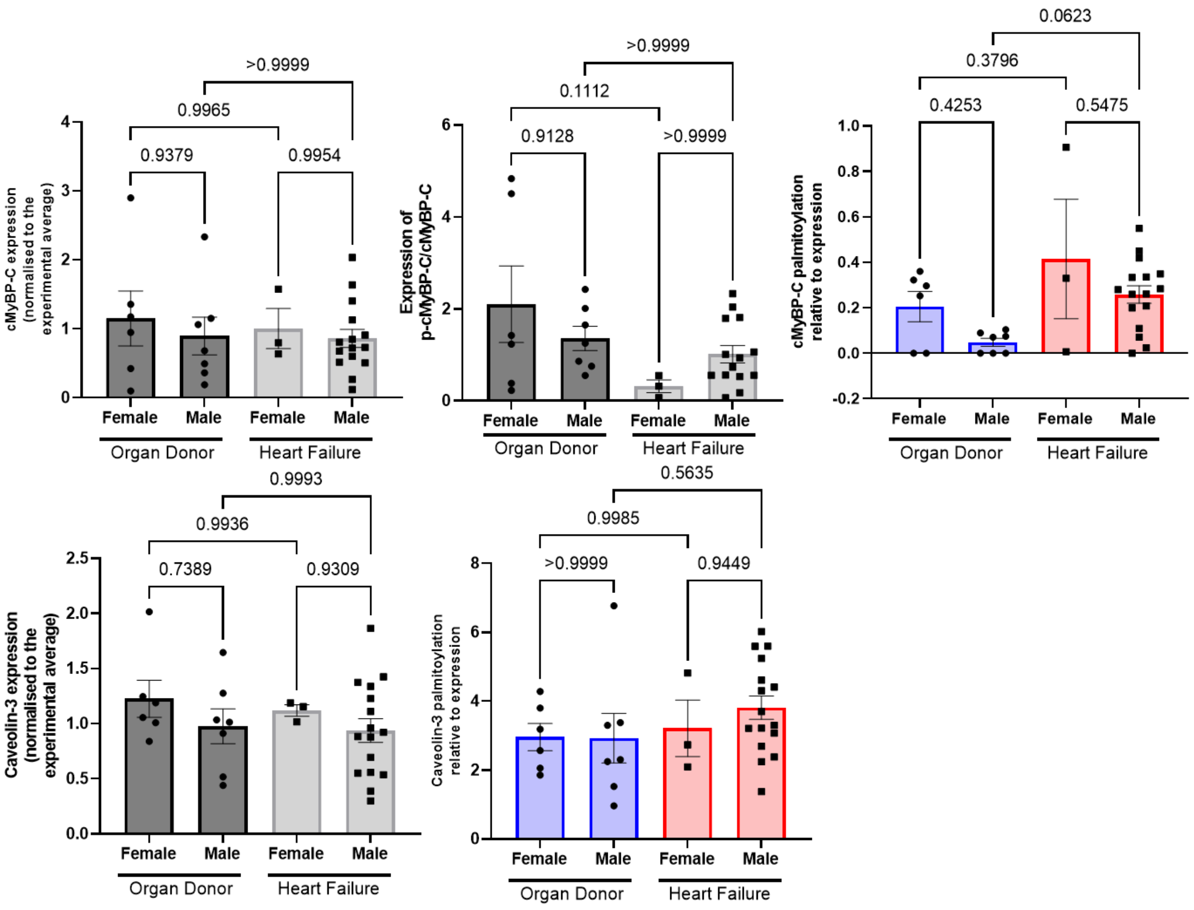


**Supplementary Figure 4. Palmitoylation of cMyBP-C in male and female organ donor and heart failure ventricular endocardium.** Palmitoylation of cMyBP-C and Caveolin-3 was determined by Acyl-Resin Assisted Capture (Acyl-RAC) in ventricular endocardium from organ donor and ischaemic heart failure, and split into groups based on sex. Palmitoylation is plotted as palmitoylated protein (HA, hydroxylamine dependent) relative to total protein (UF, unfractionated). Palmitoylation of cMyBP-C is increased in male heart failure samples (ANOVA, $p<0.05$; post-hoc comparison $p=0.06$) compared to organ donor but the same trend is not observed in females. There is no difference in overall expression however female heart failure samples may have the lowest level of p-cMyBP-C. There is no significant difference in expression or palmitoylation in Caveolin-3 between any groups. Statistical comparisons made one-way ANOVA followed by a Dunnett's post-hoc test. *$p<0.05$. Data is mean ±S.E.M.

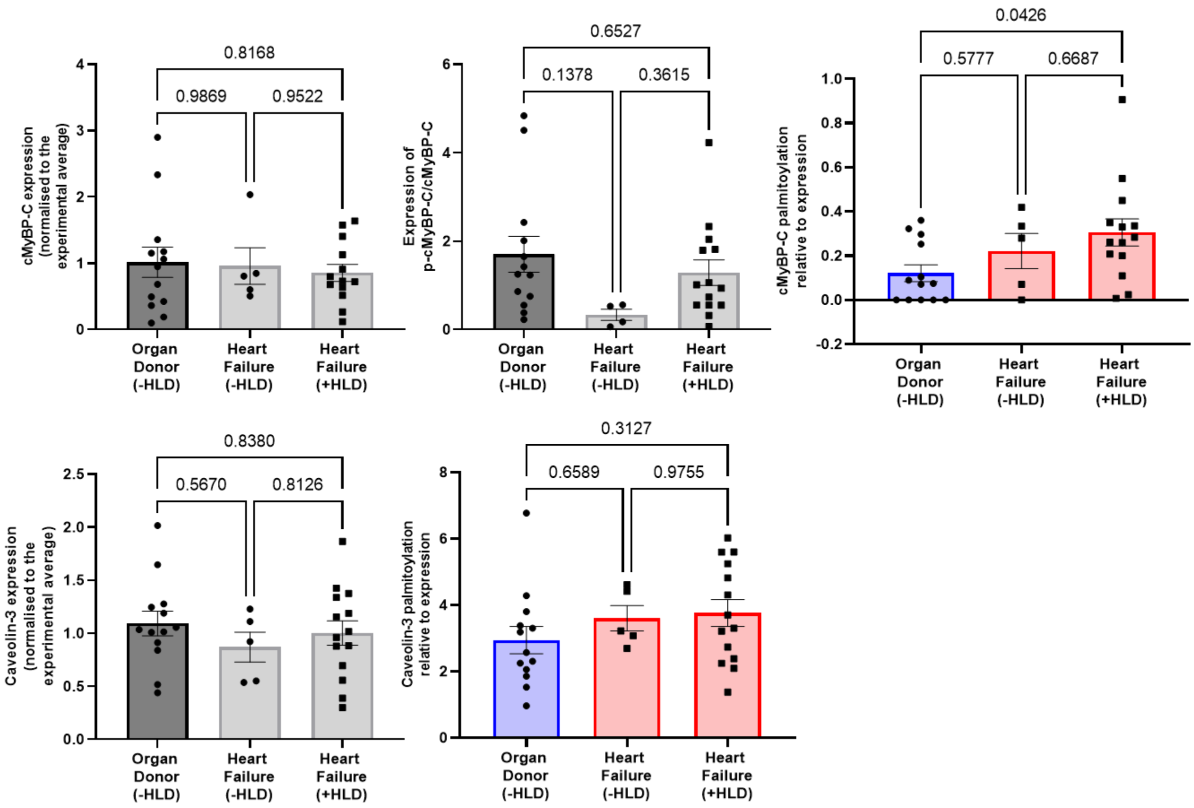


**Supplementary Figure 5. Palmitoylation of cMyBP-C in organ donor and heart failure, with and without hyperlipidemia, ventricular ednocardium.** Acyl-Resin Assisted Capture (Acyl-RAC) was used to determine the palmitoylation of cMyBP-C and Caveolin-3 in organ donor and heart failure ventricular endocardium. Total protein and level of p-cMyBP-C was also determined from the samples. Heart failure samples were then grouped into whether they classified as having hyperlipidaemia or not (all organ donors did not). Palmitoylated protein (HA, hydroxylamine dependent) was normalised to total protein (UF, unfractionated). Palmitoylation of cMyBP-C was significantly increased in heart failure samples with hyperlipidaemia, but not in those without. Whilst expression levels were unchanged between groups, samples without hyperlipidemia may have lower levels of p-cMyBP-C. Statistical comparisons made by a one-way ANOVA with a Dunnett's post hoc test. Data is mean ±S.E.M

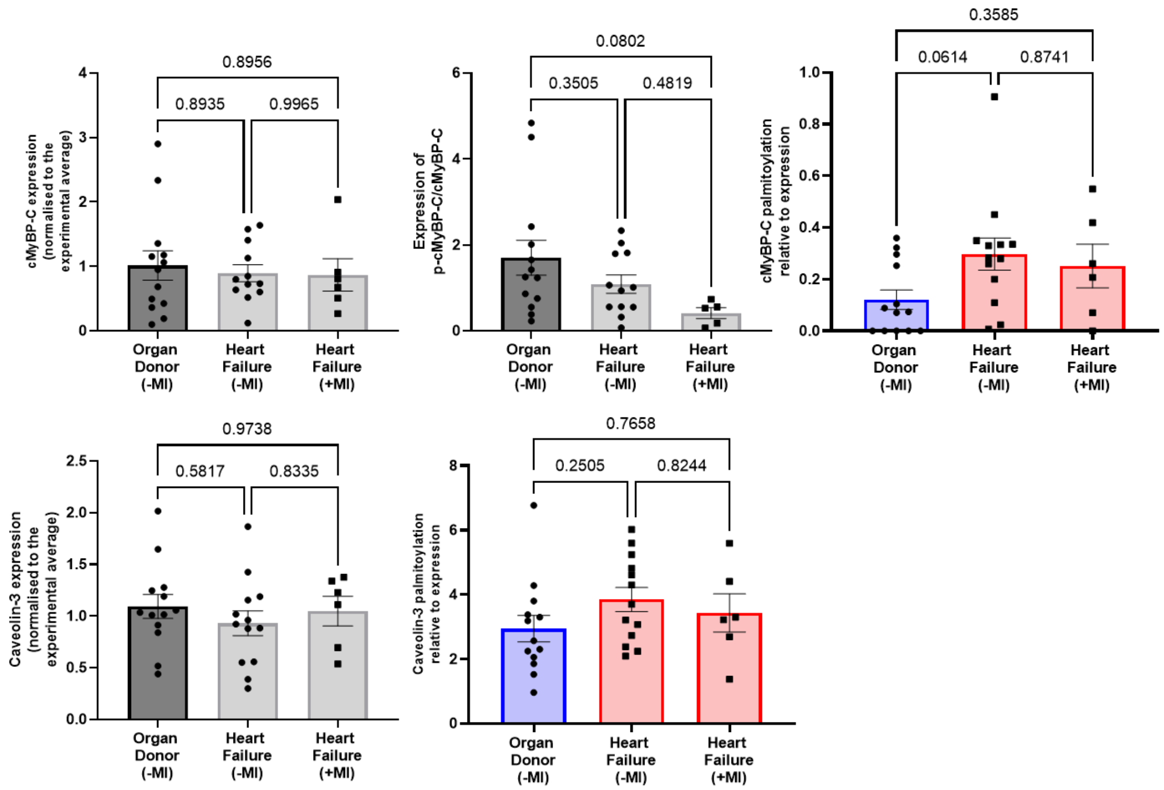


**Supplementary Figure 6. Palmitoylation of cMyBP-C in organ donor and heart failure, with and without history of myocardial infarction (MI), ventricular endocardium.** Acyl-Resin Assisted Capture (Acyl-RAC) was used to determine the palmitoylation of cMyBP-C and Caveolin-3 in organ donor and heart failure ventricular endocardium. Total protein and level of p-cMyBP-C was also determined from the samples. Heart failure samples were then grouped into whether they were listed as having a previous myocardial infarction (MI) or not (all organ donors did not). Palmitoylated protein (HA, hydroxylamine dependent) was normalised to total protein (UF, unfractionated). Palmitoylation of cMyBP-C may be most increased in samples without a history of MI. Whilst expression levels remained unchanged between groups, samples with history of myocardial infarction may have the lowest levels of p-cMyBP-C. Statistical comparisons made by a one-way ANOVA with a Dunnett's post hoc test. Data is mean ±S.E.M.

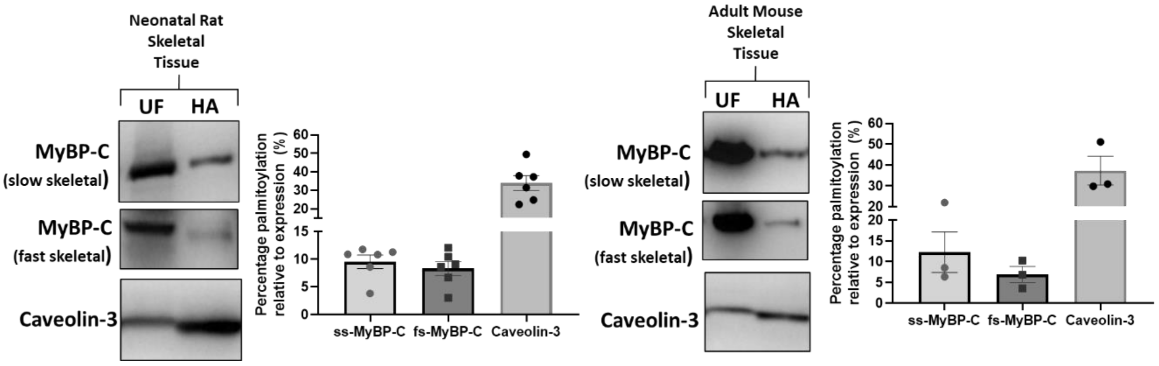


**Supplementary Figure 7. Palmitoylation of MyBP-C skeletal isoforms.** Palmitoylation of slow skeletal MyBP-C (ssMyBP-C) and fast skeletal MyBP-C (fs-MyBP-C) along with assay control Caveolin-3 was determined by Acyl-Resin Assisted Capture (Acyl-RAC) in skeletal tissue from 1-4 day old male and female neonatal rats and adult mice (male, 20 weeks). Palmitoylation (HA, hydroxylamine dependent) was normalised to total protein (UF, unfractionated) and represented as a percentage palmitoylation (%). Acyl-RAC revealed both ssMyBP-C and fsMyBP-C are palmitoylated in skeletal tissue. Data is mean ±S.E.M.
