## Supplementary Tables for "Cardiac myosin binding protein-C palmitoylation is associated with increased myofilament affinity, reduced myofilament Ca^2+^ sensitivity and is increased in ischaemic heart failure"

**Supplementary Table 1. Human heart failure and organ donor patient information.**

| Record ID | Case type | Sex | Primary diagnosis |
|---|---|---|---|
| 24713 | Organ Donor | Female | N/A |
| 2B487 | Organ Donor | Male | N/A |
| 31331 | Organ Donor | Female | N/A |
| 4B3FA | Organ Donor | Female | N/A |
| 4D931 | Organ Donor | Male | N/A |
| 5155D | Organ Donor | Male | N/A |
| 632FD | Organ Donor | Male | N/A |
| 8CB30 | Organ Donor | Female | N/A |
| B23E3 | Organ Donor | Male | N/A |
| BC90C | Organ Donor | Female | N/A |
| DOF54 | Organ Donor | Male | N/A |
| D61ZE | Organ Donor | Male | N/A |
| FC3CB | Organ Donor | Female | N/A |
| 046E | Heart Failure | Male | Ischaemic cardiomyopathy |
| 05FF7 | Heart Failure | Female | Ischaemic HFrEF |
| 14C39 | Heart Failure | Male | Ischaemic cardiomyopathy |
| 3F6DC | Heart Failure | Male | Ischaemic cardiomyopathy |
| 58545F | Heart Failure | Male | Ischaemic cardiomyopathy s/p MI |
| 6DB85 | Heart Failure | Male | HFrEF from Ischemic cardiomyopathy |
| 7CE52 | Heart Failure | Female | Ischaemic heart failure |
| 8296A | Heart Failure | Male | Ischaemic cardiomyopathy |
| 8E8D8 | Heart Failure | Male | Ischaemic cardiomyopathy |
| 97CDC | Heart Failure | Male | Ischaemic cardiomyopathy |
| 9D7E9 | Heart Failure | Male | Ischaemic cardiomyopathy |
| AF1FF | Heart Failure | Male | Chronic systolic HF |
| B8BE2 | Heart Failure | Male | Ischaemic heart failure |
| BO644 | Heart Failure | Female | Ischaemic cardiomyopathy |
| C3B57 | Heart Failure | Male | Ischaemic cardiomyopathy |
| CB8A5 | Heart Failure | Male | Ischaemic cardiomyopathy |
| DA820 | Heart Failure | Male | Ischaemic cardiomyopathy |
| EF5CB | Heart Failure | Male | Ischaemic heart failure |
| FE8E2 | Heart Failure | Male | Chronic systolic HF |

**Supplementary Table 2. Human hypertrophic cardiomyopathy and organ donor patient information.**

| Record ID | Primary Diagnosis | Mutation type |
|---|---|---|
| 1838LV | Organ Donor | |
| 1840LV | Organ Donor | |
| 1843LV | Organ Donor | |
| 1850LV | Organ Donor | |
| D6008 | Organ Donor | |
| D7040 | Organ Donor | |
| D7054 | Organ Donor | |
| HCM 113 | Hypertrophic cardiomyopathy | MYBPC3 |
| HCM 120 | Hypertrophic cardiomyopathy | MYBPC3 |
| HCM 123 | Hypertrophic cardiomyopathy | MYBPC3 |
| HCM 133 | Hypertrophic cardiomyopathy | MYBPC3 |
| HCM 169 | Hypertrophic cardiomyopathy | MYBPC3 |
| HCM 204 | Hypertrophic cardiomyopathy | MYBPC3 |
| HCM 219 | Hypertrophic cardiomyopathy | MYBPC3 |
| HCM 231 | Hypertrophic cardiomyopathy | MYL2 |
| HCM 233 | Hypertrophic cardiomyopathy | SMN |
| HCM 234 | Hypertrophic cardiomyopathy | TNNT2 |
| HCM 236 | Hypertrophic cardiomyopathy | MYH7 |
| HCM 239 | Hypertrophic cardiomyopathy | SMN |
| HCM 244 | Hypertrophic cardiomyopathy | SMN |
| HCM 255 | Hypertrophic cardiomyopathy | SMN |
| HCM 256 | Hypertrophic cardiomyopathy | MYBPC3 |
| HCM 257 | Hypertrophic cardiomyopathy | SMN |
| HCM 259 | Hypertrophic cardiomyopathy | MYH6 (VUS, class 3) |
| HCM 261 | Hypertrophic cardiomyopathy | SMN |
| HCM 263 | Hypertrophic cardiomyopathy | MYBPC3 |
| HCM 265 | Hypertrophic cardiomyopathy | MYBPC3 |
| HCM 276 | Hypertrophic cardiomyopathy | SMN |
